# Replay of Stimulus Specific Temporal Patterns during Associative Memory Formation

**DOI:** 10.1101/236059

**Authors:** Sebastian Michelmann, Howard Bowman, Simon Hanslmayr

## Abstract

Forming a memory often entails the association of recent experience with present events. This recent experience is usually an information rich and dynamic representation of the world around us. We here show that associating a static cue with a previously shown dynamic stimulus, yields a detectable, dynamic representation of this stimulus in working memory. We further implicate this representation in the decrease of low-frequency power (∼4-30 Hz) in the ongoing electroencephalogram (EEG), which is a well-known correlate of successful memory formation. The maintenance of content specific patterns in desynchronizing brain oscillations was observed in two sensory domains, i.e. in a visual and in an auditory condition. Together with previous results, these data suggest a mechanism that generalizes across domains and processes, in which the decrease in oscillatory power allows for the dynamic representation of information in ongoing brain oscillations.

## Introduction

Not everything we associate in our memory occurs at the same time. When our favorite football player is seeing the red card, for instance, we are able to bring this together with the events we just witnessed a few seconds before. Later, we are naturally able to recall all relevant information leading to the red card. In order to successfully make this association, our brain has to accomplish two things. First, it has to keep track of the past and maintain a representation of the events in the ongoing football match and second, form memories in which past events are connected to the red card. Processes during the encoding phase that will determine our ability to later remember events can be investigated with the so-called subsequent memory paradigm (Paller & Wagner, 2002; Wagner, Koutstaal, & Schacter, 1999). Subsequent memory effects refer to neural activity which distinguishes remembered from not remembered items at the time of encoding and are well documented in M/EEG and fMRI, showing involvement of cortical as well as medial temporal lobe regions (e.g. Long, Burke, & Kahana, 2014; Otten, Quayle, Akram, Ditewig, & Rugg, 2006). Concerning M/EEG power, decreases in low frequency (<40 Hz) brain dynamics have repeatedly and consistently been related to successful memory formation (Hanslmayr & Staudigl, 2014).

It has recently been proposed that cortical power decreases in the alpha/beta frequency range allow for a rich representation of memory content, since a desynchronized system has more flexibility to code information over a system of high synchrony. We call this view, the information via desynchronization framework (Hanslmayr, Staudigl, & Fellner, 2012). Confirming this idea, we have shown that sustained power decreases in the alpha band at approximately 8 Hz, contain item specific information about the remembered content, when subjects successfully replay dynamic stimuli (i.e. video and sound clips) from memory (Michelmann, Bowman, & Hanslmayr, 2016). In this study, we provided direct evidence that power decreases are involved in the representation of stimulus specific information (Hanslmayr et al., 2012). Moreover these results are well in line with numerous studies showing that perception is not continuous but rather is rhythmically sampled at a frequency of ∼7-8 Hz (Hanslmayr, Volberg, Wimber, Dalal, & Greenlee, 2013; Landau & Fries, 2012; VanRullen, Carlson, & Cavanagh, 2007). These outcomes indicate that rhythmic patterns from the perception of dynamic stimuli can reappear during internally guided retrieval processes, in the absence of the stimuli themselves. Accordingly, these prior findings also suggest the possibility that the replay of temporal patterns can be observed in a situation where dynamic stimuli have to be maintained internally in working memory.

To address this question, we here analyze the data during the encoding phase from a previous dataset (Michelmann et al., 2016). The paradigm required subjects to associate a dynamic stimulus with a static word that was used as a cue in the later retrieval phase. Importantly, during encoding the perception of the dynamic stimulus and the presentation of the word-cue was temporally separated, i.e. in every trial, one out of four dynamic stimuli was followed by a unique word-cue (Fig. 1, a-b). In a visual condition, these dynamic stimuli consisted of four short video-clips, in an auditory session four short sound clips were used. In a later retrieval block, participants were presented with the word-cue and were tested whether they remembered the associated video/sound clip.

**Figure 1:**
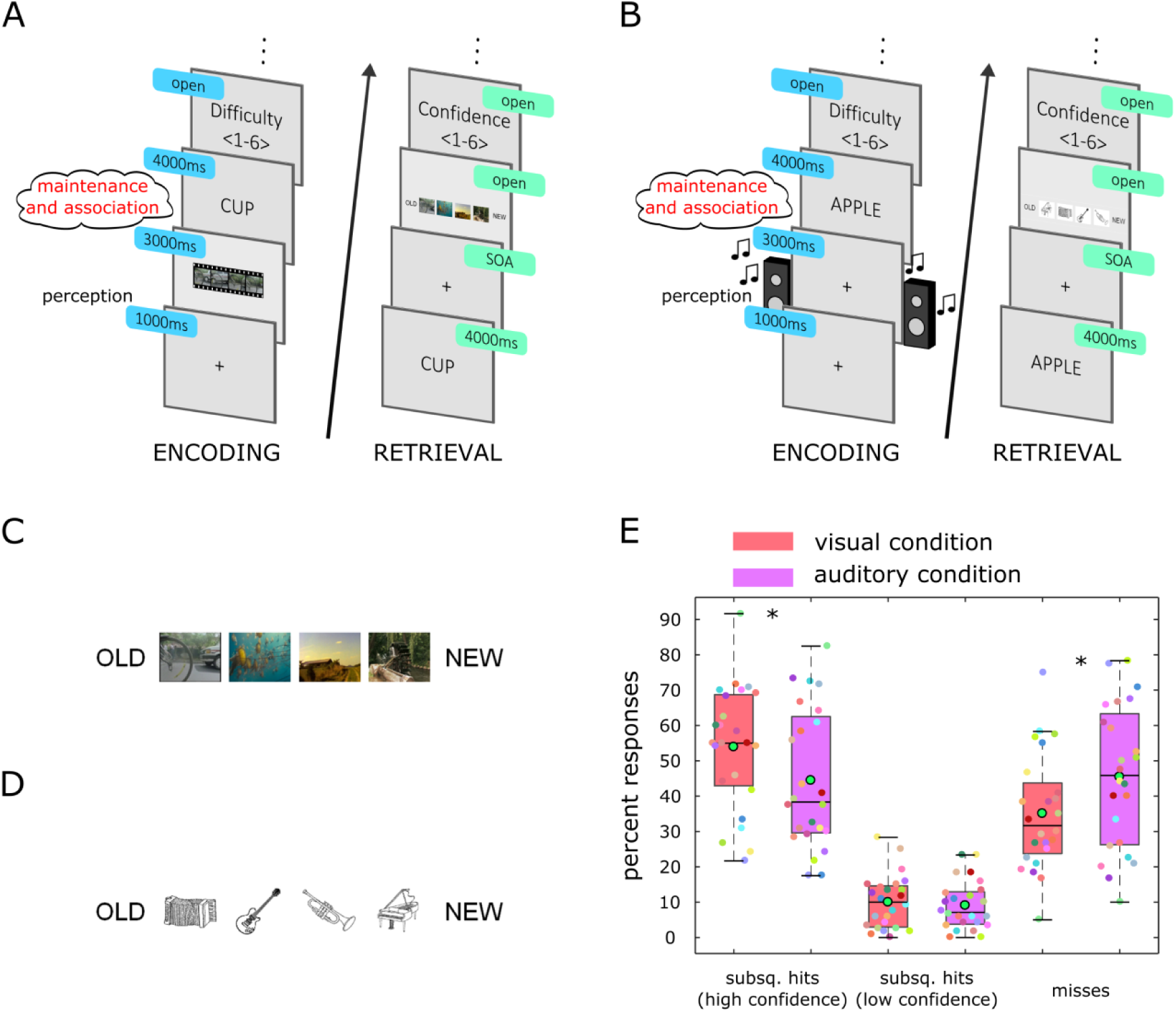
Experimental Design and behavioral results. Experimental sequence in the visual (A) and in the auditory (B) session. During encoding (A, B left), participants perceived a dynamic stimulus that played for 3 s and was then followed by a word cue. The cue was presented for 4 s and subjects had to associate the word with the dynamic stimulus they just saw. Note that during encoding, the word cue was shown after the dynamic stimulus, separating the perception interval from the association interval, therefore participants had to maintain a representation of the dynamic stimulus in working memory. Participants learned 120 associations between four repeatedly shown dynamic stimuli and 120 different words. At the end of every encoding trial, they rated the perceived difficulty of the association on a scale from 1 to 6. In the retrieval block (A, B right) they recalled the dynamic stimulus upon presentation of the word-cue. Cues from encoding were mixed with 60 new words that served as distractors. After that, they indicated the stimulus they recalled. Response options (C, D) consisted of four small screenshots of the video clips in the visual session (C) and of four small instruments, representing the sounds, in the auditory session (D). The response option “NEW” represented the distractor, and was available to indicate that the word was not presented in the encoding block, the response option “OLD” was available to indicate that subjects remembered only the word, but not it’s associate. At the end of every retrieval trial a confidence rating was collected on a scale from 1 to 6 (A, B right). (E) Behavioral performance for the associations from encoding. Hits are trials in which the correct associate was subsequently remembered (video or sound). A rating of high confidence was considered a rating > 4. Misses were defined as all trials in which the associate was later forgotten. Boxes are 25th and 75th percentiles around the median; whiskers represent minimum and maximum (disregarding outliers). Green points in black circles are arithmetic means small colored points are single subject values. * p < 0.05

We hypothesize that, in order to associate the word-cue with the dynamic stimulus, subjects maintain (i.e. replay) a sensory representation of the dynamic stimulus in working memory, which is why we refer to this phase as the maintenance phase. Using temporal pattern similarity analysis, we should therefore be able to detect the replay of these patterns during the maintenance phase, i.e. when the association between a word and the sound/movie is formed. In accordance with the information via desynchronization framework, we should observe stronger decreases for later remembered versus later not remembered items. This subsequent memory effect should be most evident in the frequency band that codes for the representation of the dynamic stimulus in working memory, i.e. 8 Hz as per our previous findings. Moreover, if power decreases enable a richer representation of the perceptual content, we should already observe stronger broad power decreases for later remembered compared to later not remembered stimuli during the perception of the dynamic stimulus.

## Materials and Methods

### Participants

24 healthy, right-handed subjects (18 female and 6 male) participated in this study. 7 further participants were tested, or partly tested, but could not be analyzed due to poor memory performance (N=2), misunderstanding of instructions (N=2), and poor quality of EEG-recording and technical failure (N=3). All participants had normal or corrected-to normal vision. The average age of the sample was 23.38 (s.d. = 3.08) years. Participants were native English speakers (20), bilingual speakers (2) or had lived for more than 8 years in the UK (2). Ethical approval was granted by the University of Birmingham Research Ethics Committee, complying with the Declaration of Helsinki. Participants provided informed consent and were given a financial compensation of 24£ or course-credit for participating in the study.

### Material and experimental set up

The cues amounted to 360 words that were downloaded from the MRC Psycholinguistic Database (Coltheart, 1981). Stimulus material consisted of 4 video clips and 4 sound clips in the visual and auditory session respectively. All clips were 3 seconds long; videos showed colored neutral sceneries with an inherent temporal dynamic, sounds were short musical samples, each played by a distinct instrument. In both sessions, a clip was associated with 30 different words. 60 words were reserved for the distractor trials and 12 additional words were used for instruction and practice of the task. For presentation, words were assigned to the clips or to distractors in a pseudorandom procedure, such that they were balanced for Kucera-Francis written frequency (mean = 23.41, s.d. = 11.21), concreteness (mean = 571, s.d. = 36), imageability (mean = 563.7 s.d. = 43.86), number of syllables (mean = 1.55, s.d. = 0.61) and number of letters (mean = 5.39, s.d. = 1.24). Furthermore lists were balanced for word-frequencies taken from SUBTLEXus (Brysbaert & New, 2009). Specifically, “Subtlwf” was employed (mean = 20.67, s.d. = 27.16). The order of presentation was also randomized, assuring that neither the clips and their associates, nor distractor words were presented more than 3 times in a row or in temporal clusters. The presentation of visual content was realized on a 15.6 inch CRT-monitor (Taxan ergovision 735 TC0 99) at a distance of approximately 50 centimeters from the subjects eyes. The monitor refreshed at a rate of 75 Hz. On a screen size of 1280 × 1024 pixels, the video clips appeared in the dimension of 360 pixels in width and 288 pixels in height. ‘Arial’ was chosen as the general text-font, but font-size was larger during presentation of word-cues (48) than during instructions (26). In order to reduce the contrast, white text (rgb: 255, 255, 255) was presented against a grey background (rgb: 128, 128, 128). Auditory stimuli were presented using a speaker system (SONY SRS-SP1000). The 2 speakers were positioned at a distance of approximately 1.5 meters in front of the subject with 60 centimeters of distance between the speakers.

### Procedure

Upon informed consent and after being set up with the EEG-system, participants were presented with the instructions on the screen. Half of the subjects started with the auditory session, the others were assigned to undertake the visual task first. Both sessions consisted of a learning block, a distractor block and a test block. The sessions were identical in terms of instructions and timing and differed only in the stimulus material that was used. During instruction, the stimulus material was first presented for familiarization and then used in combination with the example words to practice the task. Instructions and practice rounds were completed in both sessions.

As a way to enhance memory performance, participants were encouraged to use memory strategies. The suggestion was to imagine the word in a vivid interaction with the material content, yet the choice of strategy remained with the subject. In the learning block, 120 clip-word sequences were presented. Each sequence started with a fixation cross that was presented in the center of the screen for 1 second, and then the video-clip played for 3 seconds. In the auditory condition, the fixation cross stayed on the screen and the sound-clip played for 3 seconds. Immediately after the clip, a word cue was presented in the center for 4 seconds, giving the subject time to learn the association. After that, an instruction requested participants to subjectively rate on a 6 point scale how easy the association between the clip and the word was. After a press on the space bar, this scale was shown. Equidistant categories were anchored with the labels “very easy” and “very hard”; those labels were displayed at both ends above the scale. Participants used six response buttons to rate the current association (see: Fig. 1).

In the distractor block, subjects engaged in a short unrelated working memory task, namely they counted down in steps of 13, beginning from 408 or 402 respectively. After 1 minute the distractor task ended. Following a short self-paced break, subjects refreshed the instructions on the retrieval block.

In this retrieval block, either a cue or a distractor was presented upon a button press on the space bar. Subjects were instructed to try to vividly replay the content of the corresponding video-clip or sound-clip in their mind upon presentation of the cue. The word stayed on the screen for 4 seconds, giving the subject the opportunity to replay the memory. Finally, a fixation cross was presented for a varying time window between 250 and 750 milliseconds to account for movement and preparatory artifacts, before the response scale appeared on the screen.

The response-scale consisted of 6 options. 4 small screen shots of the videos or 4 black and white pictures of the featured instruments were presented in equidistant small squares of 30x30 pixels. Additionally, the options “new” and “old” were displayed in the form of text at the most left and most right position of the scale (see: Fig. 1 C-D). Subjects could now either indicate the target (video/sound) they just replayed, by pressing the button corresponding to that clip. Instead, subjects could also indicate that the word was a distractor by pressing the button corresponding to the option “new”, or they would simply indicate that they remembered the word, but could not remember the clip it was associated with. In this last scenario, subjects would press the button corresponding to "old”. The positions of “old” and “new” at the end of the scale, as well as the permutation of the 4 target positions in the middle of the scale, were counterbalanced across participants. Finally, after making a decision, a further six point rating scale was presented on which subjects could rate the confidence in their response. Again a scale with equidistant categories was presented ranging from “guess” to “very sure”. An additional possibility was to press “F2” in case of an accidental wrong button press. In this case, the whole trial was discarded from analysis. Following the retrieval block, individual electrode positions were logged allowing for a break of approximately 30 minutes before beginning the second session.

### Data Collection

The recording of behavioral responses and the presentation of instructions and stimuli were realized using Psychophysics Toolbox Version 3 (Brainard, 1997) with MATLAB 2014b (MathWorks) running under Windows 7, 64 Bit version on a desktop computer. Response buttons were “s, d, f, j, k, l” on a standard “QWERTY” layout. Buttons were highlighted and corresponded spatially to the response options on the screen, so participants did not have to memorize the keys. To this end, the shape of corresponding fingers was also displayed under the scale. To proceed, participants used the space bar during the experiment. Physiological responses were measured with 128 sintered Ag/AgCl active electrodes, using a BioSemi Active-Two amplifier, the signal was recorded at 1024 Hz sampling rate on a second computer via ActiView recording software, provided by the manufacturer (BioSemi, Amsterdam, Netherlands). Electrode positions were logged with a Polhemus FASTRAK device (Colchester, VT, USA) in combination with Brainstorm (Tadel, Baillet, Mosher, Pantazis, & Leahy, 2011) implemented in MATLAB.

### Preprocessing

The data was preprocessed using the Fieldtrip toolbox for EEG/MEG-analysis (Oostenveld, Fries, Maris, & Schoffelen, 2011). Data was cut into trial-segments from 2.5 seconds prestimulus to 7 seconds after the onset of the dynamic stimulus. The linear trend was removed from each trial and a baseline correction was applied based on the whole trial. Trials were then downsampled to 512 Hz and a band-stop filter was applied at 48-52, 58-62, 98-102 and 118-122 Hz to reduce line noise at 50 Hz and noise at 60 Hz; additionally a low-pass filter at 140 Hz was applied. After visual inspection for coarse artifacts, an independent component analysis was computed. Eye-blink artifacts and eventual heartbeat/pulse artifacts were removed, bad channels were interpolated and the data was referenced to average. Finally the data was inspected visually and trials that still contained artifacts were removed manually.

### Behavioral analysis

For behavioral analysis, correct trials were defined as those in which the target was correctly identified. The confidence rating of the response was considered as high if a rating of 5 or 6 was selected. Misses were defined as trials in which a cue-word was incorrectly identified as a new word, the wrong clip was selected, or the response “old” was given to indicate recognition of the word without remembering the target video or sound it was associated with.

### Power analysis

Oscillatory power was determined by multiplying the Fourier-transformed data with a complex Morlet wavelet of 6 cycles. Raw power was defined as the squared amplitude of the complex Fourier spectrum and estimated for every 4^th^ sampling point (i.e. sampling rate of 128 Hz). For the contrast of subsequent hits and subsequent misses, a baseline was computed as the average power between −1 and 7 seconds of all trials within the contrast (Long et al., 2014). Every trial was then normalized by subtracting the baseline and subsequently dividing by the baseline (activity_tf_ – baseline_f_)/baseline_f_, where t indexes time and f indexes frequency. The relative power was calculated for all frequencies between 2 and 30 Hz.

### Phase pattern analysis during perception and maintenance

While participants learned the associations in the encoding block, they repeatedly perceived (saw/heard) the same dynamic stimulus. Content-specific properties could consequently be identified if they were shared by trials of the same content but not by trials of a different content. Hence, content-specific phase during perception was assessed by contrasting the phase similarity between pairs of trials in which the same content was presented, with the phase similarity of an equal number of trial pairs that were of different content. For each pair of trials, the cosine of the absolute angular distance was then computed and finally averaged across all (same or different) combinations [29]. The average similarity value for same and different combinations was subjected to statistical testing across subjects at every time point, at every electrode and in every frequency of interest; this contrast embodies content specific phase patterns during perception.

Participants also repeatedly associated the same dynamic stimulus (one of four videos/sounds) with a different word cue. Therefore the temporal pattern during perception of the dynamic stimulus could be compared to the temporal pattern in different trials in which subjects maintained the same dynamic stimulus in working memory. Notably, excluding within trial combinations eliminates the potential confound of temporal autocorrelation. Likewise the temporal pattern during perception could be compared to trials in which subjects maintained a different dynamic stimulus in working memory.

In this way, the phase similarity between combinations of same content (e.g. perceiving content 1, maintaining content 1) was contrasted with the phase similarity between trials of different content (e.g. perceiving content 4, maintaining content 2). This contrast reveals phase patterns that are specific to the dynamic stimulus which subjects associated with the cue.

To maximize the signal to noise ratio, the following restrictions were applied: The tested frequency was 8 Hertz, following our previous results and hypotheses (Michelmann et al., 2016); A time-window during perception was centered on the cluster in which phase patterns were most reliably content specific during encoding (i.e. the cluster with the lowest p-value) and subsequently used in a sliding window approach in order to detect content specific patterns.

Phase similarity between two windows was then assessed with the Single-trial Phase Locking Value (S-PLV) (Lachaux et al., 2000; Mormann, Lehnertz, David, & E. Elger, 2000). This measure defines similarity between two windows (x and y) as the constancy of phase angle difference over time, where n denotes the width of the window and *φ* is the phase:

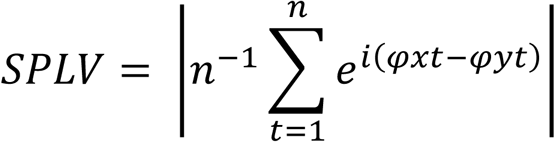

S-PLV assesses the phase coherence between two time windows and has the advantage of increased robustness for noisy data at the expense of temporal resolution.

(Lachaux et al., 2000) suggest to compute the S-PLV over 6-10 cycles of a frequency for a good signal to noise ratio, for our purposes, S-PLV was applied to a time window of 8 cycles, which resulted in a 1 second window for 8 Hertz. Phase values were extracted by multiplying the Fourier-transformed data with a complex Morlet Wavelet of 6 cycles. Phase-values were then downsampled to 64 Hz. The similarity measure was computed for every pair of trials in the combinations of same content and in the combinations of different content. Importantly, a sliding window approach was used to account for the non-time-locked nature of the data (temporal patterns could be present anywhere in the maintenance interval). This resulted in a time course of similarity for the combinations of same and of different content.

The difference in this similarity was first averaged across the whole maintenance episode (between 3500 and 7000ms) and then statistically tested across subjects with a random permutation procedure based on clusters of summed t-values across electrodes (Maris & Oostenveld, 2007). In a second test, the time courses at every electrode were compared with a series of t-tests and subsequently tested with a cluster-based random permutation procedure, where clusters were summed across electrodes and time (see also: statistical analyses, below). Additionally, a control frequency was tested, namely 6 Hz, based on the results from the power analysis. Time windows were defined accordingly for this frequency as 8 cycles around the center of the most reliable cluster during perception.

### Statistical analyses

#### Behavioral performance

Behavioral results were compared between the auditory and visual condition with a series of paired t-tests. P-values were compared against a Bonferroni-corrected threshold (Bland & Altman, 1995), however no specific hypothesis was tested.

#### Decreases in power

To test for differences in baseline corrected power, a paired t-test was first computed for every time point and frequency at every channel. For multiple-comparison-correction, a random permutation procedure was applied (Maris & Oostenveld, 2007). This procedure sums up neighboring t-values above a cluster forming threshold and compares the resulting clusters’ sizes to the distribution of the maximal cluster sums that are derived, when condition labels are randomly swapped with the Monte-Carlo method. The minimum number of neighboring channels to be considered a cluster was specified with 3, which attenuates the impact of spatially high frequency noise; neighboring electrodes were derived via the triangulation method of the Fieldtrip toolbox (http://www.fieldtriptoolbox.org/). The clusters were summed across time, frequency and channels, then labels were permuted 1000 times; thresholding of the clusters as well as the testing of the null hypothesis was addressed with a threshold for single-sided testing (alpha level of 0.05). To identify frequencies with a reliable power difference, a paired samples t-test was computed for every frequency on the average power difference across all channels and across the whole time window of interest.

#### Phase similarity during perception of the dynamic stimulus

Phase similarity during perception was tested in the same way as power. A series of paired t-tests was computed to contrast the average similarity of combinations of same content with the average similarity of combinations of different content. T-values for every frequency band, electrode and time point were then corrected for multiple comparisons in an unrestricted cluster-based permutation approach. The cluster permutation compared again the sums of t-values across frequency, electrodes and time against the distribution of these clusters derived via the Monte-Carlo method. Later, the frequency 8 Hertz was tested separately with the same cluster permutation in order to identify a temporospatial cluster, in which 8 Hertz phase could differentiate content particularly well.

#### Phase similarity between perception and maintenance

The similarity between the time-window during perception and the maintenance-episode was tested for differences between combinations of same and combinations of different content. As mentioned above, in a first step, the average difference between 3.5 and 7 seconds was contrasted with a paired t-test on every electrode to test for a general effect. For multiple comparisons correction, again, 1000 permutations were drawn. Observed clusters of those t-values that exceeded the critical threshold were summed across neighboring electrodes and were tested against the distribution of sums under random permutation of conditions. In a second step, a paired t-test was computed for every electrode and time point during the maintenance interval and differences were again tested with a cluster based permutation approach. Now clusters were formed by summation of the thresholded t-values across electrodes and time and compared against the distribution of these clusters for 1000 random permutations.

## Results

### Behavioral performance

In the visual session, participants remembered on average 53.92% (standard deviation [s.d.] = 17.56%) of the video clips with high confidence (rating > 4), and they further remembered 9.97% (s.d. = 7.62%) of the clips with low confidence (Fig. 1 E). In the auditory session, 44.44% (s.d. = 19.8%) of the audio clips were subsequently remembered with high confidence, which was significantly less than in the visual condition (*t*_23_ = −2.81, *p* < 0.01). An additional 9.06% (s.d. = 6.9%) of the audio clips were remembered with low confidence. In accordance, the number of subsequent misses was significantly lower in the visual session (mean 35.07%, s.d. = 16.43%) than in the auditory session (45.45%, s.d. = 20.27%, *t*_23_ = −3.33, *p* < 0.01).

### Successful memory encoding is associated with low frequency power decreases in the visual and auditory condition

To find correlates of successful memory encoding, the oscillatory power between subsequently remembered (hits) and subsequently not remembered (misses) items was compared. Specifically, we contrasted trials for which associations were subsequently remembered with high confidence, with trials in which the associations were subsequently not remembered correctly. In this analysis, only those datasets were used, in which a minimum of 15 trials remained for hits or misses after preprocessing (N=18). Two crucial episodes for successful memory encoding were tested separately: (i) the time interval in which the dynamic stimulus was actually perceived (0 to 3 seconds) and (ii) the maintenance interval (3 to 7 seconds), in which the memory formation would be expected to have taken place. In the time interval from 0 to 3 seconds, a small cluster of power decreases was associated with successful memory in the visual condition; it displayed a trend towards significance (*p* < 0.07, Fig. 2A, left). Likewise, in the auditory condition a similar cluster of power decreases appeared (*p* = 0.047, Fig. 2B, left).

**Figure 2:**
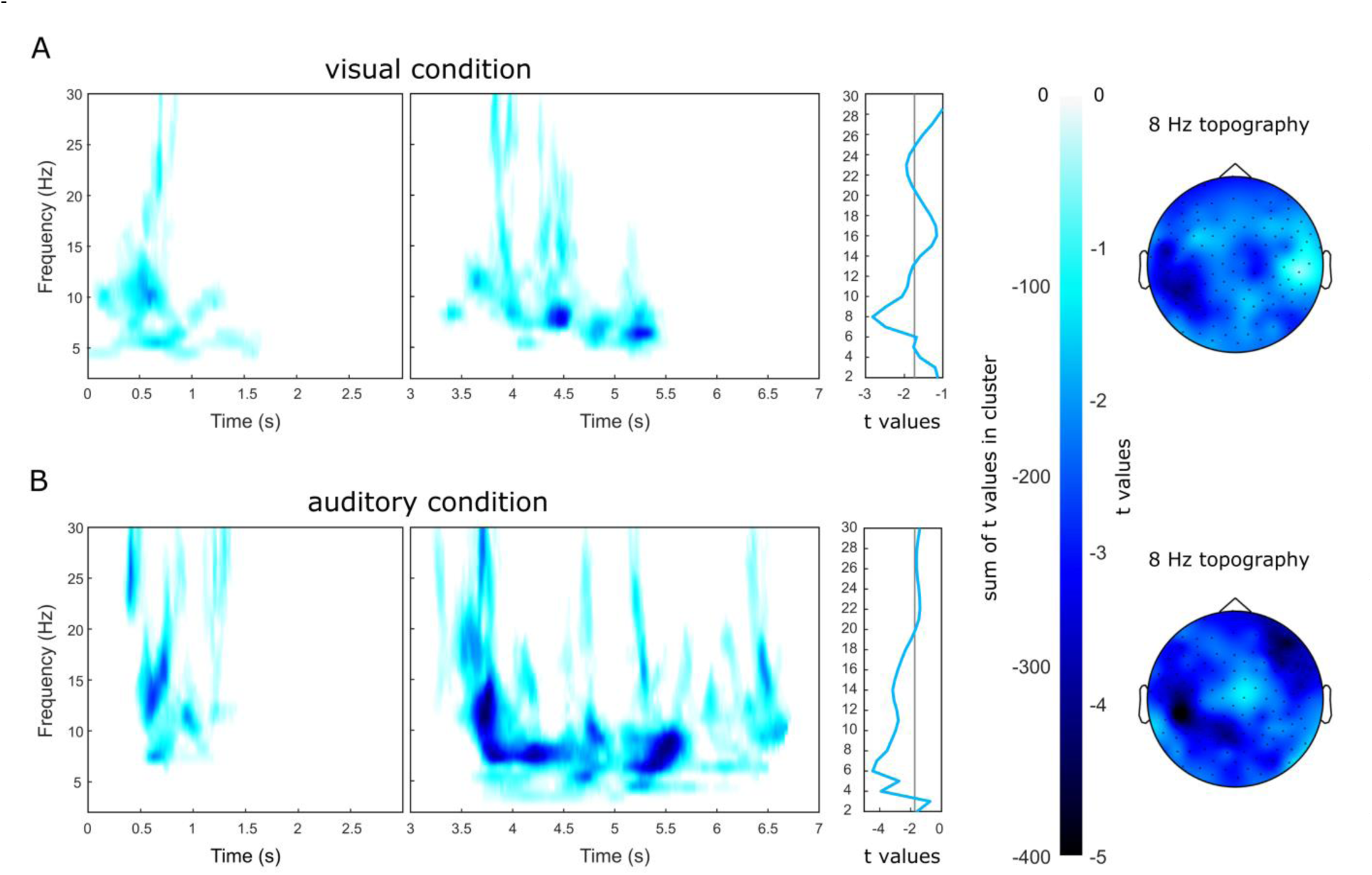
Subsequent memory effects in oscillatory power. Successful memory encoding was associated with broad power decreases in the lower frequencies (<30 Hz) in the visual (A) and in the auditory (B) condition. In the first 3 seconds of a trial, when subjects perceived the dynamic stimuli, clusters of broad power decreases in the visual (A, left) and auditory (B, left) condition displayed a trend towards significance already during this interval. During the association with the word-cue (between 3 and 7 seconds within each trial) broad clusters of significant power decreases emerged in both conditions (A, B, middle). Time frequency plots show the sum of t-values across the clusters (A and B, left and middle panels). The t-value of average power difference across electrodes and time between 3 and 7 seconds is plotted to the right of the middle panels. Topographies of power decreases are plotted on the right as maps of t-values derived from the average power decreases between 3 and 7 seconds.

During the maintenance interval (3 to 7 seconds), substantially reduced power in the lower frequencies (<30 Hz) was observed for subsequent hits compared to subsequent misses (Fig. 2, middle) in both conditions. In the visual condition, a broad cluster emerged where power was significantly lower when tested against random permutations (*p* = 0.031, Fig. 2A, middle). Likewise a broad cluster of significant power decreases appeared in the maintenance interval of the auditory condition (*p* < 0.003, Fig. 2B, middle).

To identify frequencies that robustly exhibited lower oscillatory power for successful memory encoding, the power during the maintenance interval was averaged across all electrodes and time points and differences were subjected to a t-test. Following our previous results (Michelmann et al., 2016), we expected the strongest power decreases in both conditions to peak at 8 Hz. Indeed, a clear peak at 8 Hz was observed in the visual condition (*t*_17_= −2.82, *p* < 0.01, Fig. 2A, middle). In the auditory condition, however, a peak was observed at 6 Hz (*t*_17_ = - 4.45, *p* < 0.001, Fig. 2B, middle), yet power decreases also extended to 8 Hz (*t*_17_ = −3.53, *p* = 0.001).

For the visual condition, the power decreases at 8 Hz displayed a broad topography with a parietal maximum over the left hemisphere (Fig. 2A, right). Decreases in 8 Hz power were similarly broadly distributed in the auditory condition, with maxima over left parietal and right frontal regions (Fig. 2B, right).

Together, these results confirm the fundamental role of decreases in low frequency oscillatory power for the successful formation of memory.

### Temporal patterns are content specific during perception and can be detected during maintenance

The detection of content specific temporal patterns during the maintenance period necessitates that the dynamic stimuli themselves elicit temporally distinct neural responses. To address this, we first compared the pairwise phase consistency (PPC) (Vinck, van Wingerden, Womelsdorf, Fries, & Pennartz, 2010) between trials in which the same dynamic stimulus was perceived with the PPC between trials of different content. Oscillatory phase of the neural responses was specific to the dynamic stimuli in two broad clusters in the visual (*p* < 0.001, *p* = 0.003, Fig 3C) and one broad cluster in the auditory condition (*p* < 0.001, Fig 3G), confirming prior reports that the content of dynamic stimuli is tracked by the phase of low frequency oscillations (Ng, Logothetis, & Kayser, 2013). Vitally, both clusters included 8 Hz which was the oscillation for which we hypothesized to detect the reappearance of temporal patterns in the maintenance period.

**Figure 3:**
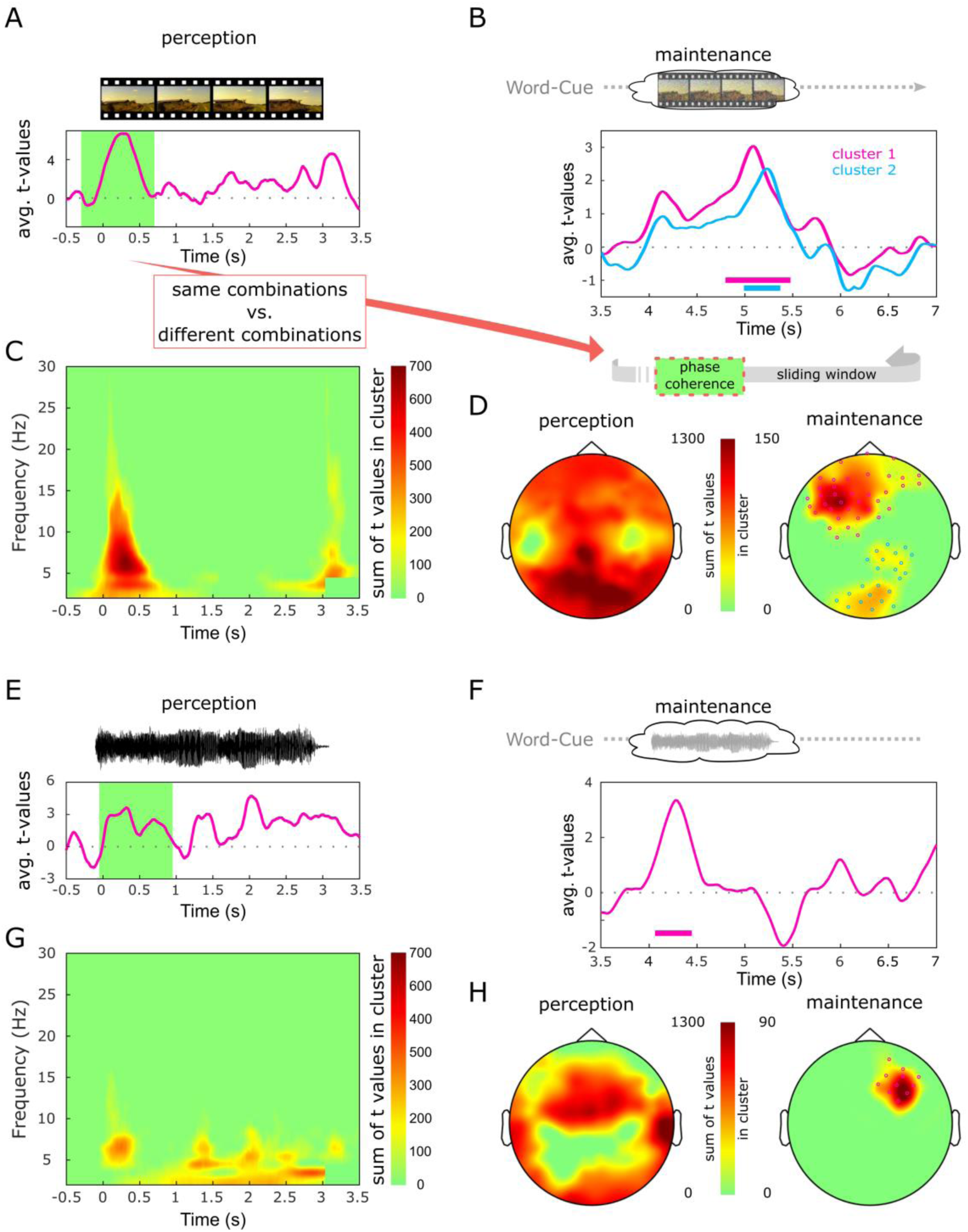
Content specificity of oscillatory phase during perception and maintenance of a dynamic stimulus. Oscillatory phase distinguishes between content during perception (left) and can be detected again during maintenance (right). During perception, pairwise phase consistency between trial-combinations of same content and combinations of different content was contrasted with t-tests (A, C, E, and G). (C) shows these t-values for every time and frequency bin in the visual condition, (G) displays the auditory condition. A horizontal slice through these time frequency plots is represented in (A) and (E) for the frequency of 8 Hz. The green window denotes the time window that was selected in order to detect content specific maintenance in the subsequent time interval, where only a static word-cue was presented on the screen. A measure of phase coherence over time (S-PLV) was computed between the selected window and every time point during the maintenance period (A-B).Importantly this similarity was never computed within trials, in order to balance temporal autocorrelations. The result from this sliding window approach was tested with a series of t-tests contrasting same and different content-combinations (e.g. watching movie 1, maintaining movie 1 vs. watching movie 2, maintaining movie 4). (B) and (F) show the time-courses of t-values within the clusters of significant differences in the visual (B) and auditory (F) condition. Horizontal bars denote the time interval in which clusters emerged with significant differences. (D) and (H) display the sum of t-values across the selected time window during perception (D, H, left) and across time within clusters, during maintenance (D, H, right).

We now identified periods during perception in which the time courses at 8 Hz were maximally content specific by restricting the statistical test to 8 Hz only and selecting the cluster in which content could most reliably be differentiated during perception (i.e., the cluster with the lowest p-value). In the visual condition, this cluster extended from −152 ms to 564 ms (*p* < 0.001). Note that post-stimulus effects are smeared temporally into the prestimulus interval because of the wavelet decomposition. The most reliable cluster of content specificity in the auditory condition extended from 22 ms to 871 ms (*p* = 0.002). A further cluster in the visual condition was observed between 2,650 ms and 3,300 ms (*p* = 0.016). In the auditory condition further clusters emerged between 1,818 ms and 2,627 ms (*p* = 0.003) and between 1,203 ms and 1,504 ms (*p* = 0.047) indicating that in both modalities early and later time windows showed content specific temporal patterns.

For the 8 Hz oscillation, a 1-second wide window was now centered on the cluster that most reliably distinguished content during perception (i.e. at 206 ms in the visual condition and 446 ms in the auditory condition, Fig 3 A, E). In a sliding window approach, a measure of phase coherence (S-PLV (Lachaux et al., 2000)) was then computed between this window and every 1-second-wide window between 3 and 7 seconds during the maintenance period (see Fig. 3 A-B). For practical reasons, at the end of the trial the window was slid out back into the pre-stimulus interval (zero padding could be an alternative but more intricate approach). This time course of similarity (phase coherence) was now computed for trial-combinations comprising perception and maintenance of the same stimulus and for trial-combinations of perception and maintenance of different content. Importantly, the combinations of same content were never built within a trial, assuring a balancing of temporal autocorrelation between same and different combinations. In a first test, we subjected the average similarity across time to a t-test, contrasting same and different combinations at every electrode. A cluster-based permutation revealed a significant cluster in the visual condition (*p* < 0.001), but not in the auditory condition. In a follow-up test, we repeated the t-test for every time-point at every electrode and summed clusters across time and electrodes. A permutation test revealed 2 clusters of significant differences in the visual condition (*p* < 0.001, *p* = 0.035, Fig. 3 B, D). The first cluster was located over left-frontal regions and extended from 4.8 to 5.41 seconds after stimulus onset (i.e. 1.8 to 2.41 seconds after the start of the maintenance phase). The second cluster was located over parietal and occipital areas, extending from 4.97 to 5.34 seconds (1.97 to 2.34 seconds of the maintenance phase, Fig. 3 D right). We applied the same approach to the auditory condition; a cluster (*p* = 0.047) emerged over right-frontal regions extending from 4.11 to 4.44 seconds after stimulus onset (1.11 to 1.44 seconds of the maintenance phase), even though strictly interpreted, this cluster does not exceed a corrected alpha threshold (Fig. 3 F, H right). Finally we also tested the frequency of 6 Hz, which showed the most reliable power decrease in the auditory condition, however no effects were found.

## Discussion

For most of the memories that we form during the day, we rely on rich and dynamic ongoing representations of the world around us. At a later point, we then associate these representations with distinct events. Both of these properties of our natural experience are rarely captured in experiments that investigate episodic memory. First, most studies use non-information rich stimuli to study memory, like words or pictures, and second material for association is usually presented simultaneously.

In this study, we used a memory task that can mimic memory in a more naturalistic scenario: an ongoing representation of an information rich, dynamic stimulus is maintained in working memory, in order to be associated with a subsequent event. In one session, subjects repeatedly watched one out of four short video clips, which was immediately followed by a unique word-cue. In a second session, subjects listened to one out of four sound clips, which they subsequently associated with a cue (Fig. 1). In order to form an association, participants had to maintain a representation of the video/sound clip in working memory.

Investigating the correlates of subsequent memory, we found broad and sustained decreases in ongoing oscillatory power to be associated with successful memory formation. These power decreases were particularly strong while subjects maintained dynamic representations in working memory, namely while they formed the association. Importantly, we found that these power decreases carried stimulus specific information in their temporal pattern of activity. Specifically, the phase of an 8 Hz frequency, which we previously linked to content representation (Michelmann et al., 2016) and where power decreases were strongest in the visual condition, was modulated in a stimulus specific way.

These results form part of converging evidence for a general mechanism, in which desynchronization of brain oscillations in the cortex, indicated by power decreases, allows for the rich representation of information (Hanslmayr et al., 2012). Specifically, the decrease in oscillatory strength, which also signifies a release from inhibition (Haegens, Nacher, Luna, Romo, & Jensen, 2011; Klimesch, Sauseng, & Hanslmayr, 2007), renders the oscillation less stationary, i.e. less predictable. In mathematical terms, this decrease of predictability means an increase in the amount of information that can be coded (Hanslmayr et al., 2012; Shannon & Weaver, 1949). When we previously observed this mechanism during episodic memory reinstatement, oscillatory patterns were localized in sensory-specific areas (Michelmann et al., 2016). In contrast, the pattern maintenance observed in this analysis displayed a different, i.e. more frontal topography, which is suggestive of working memory processes (e.g. Goldman-Rakic, 1995). The generalization of this desynchronization-mechanism across different processes is further complemented by its generalization across modalities; namely, in this study as well as in previous results, we observed oscillatory patterns in desynchronizing brain dynamics for visual and auditory stimuli.

Finally, the frequency band of 7-8 Hz has been previously implicated in the rhythmic sampling of perceptual content (Hanslmayr et al., 2013; Landau & Fries, 2012; VanRullen et al., 2007). These studies integrate well with our findings and suggest that the 8 Hz frequency temporally organizes the representations of stimulus specific information during perception, episodic memory reinstatement and working memory maintenance and that decreases in oscillatory power allow these temporal patterns to resurface.

Our results moreover inform current debates about the neural mechanisms underlying working memory. While some studies have previously shown that content specific activity patterns can be decoded during working memory maintenance (Fuentemilla, Penny, Cashdollar, Bunzeck, & Düzel, 2010; Jafarpour, Penny, Barnes, Knight, & Duzel, 2017), other studies suggest that representations in working memory may not always be maintained online, but rather latently stored in synaptic weights or even via more complex mechanisms (see: Stokes, 2015, for review). Those representations can then reemerge when they become task relevant, or they can be evoked experimentally by either ‘pinging’ them with unspecific input (Wolff, Jochim, Akyürek, & Stokes, 2017) or by stimulating transcranially with a magnetic pulse (Rose et al., 2016). Hence, an important insight from the here presented study is that in order to form an association with a previously shown dynamic stimulus, a representation of that stimulus is maintained online in working memory.

The method that we used in order to observe these stimulus patterns was specifically tailored to the detection of patterns that are dynamic in nature. This is very relevant for studies that investigate working memory maintenance because patterns that are involved in the online maintenance of representations in Prefrontal Cortex and Parietal Cortex of nonhuman primates, have been found to be highly dynamic (Crowe, Averbeck, & Chafee, 2010; Meyers, Freedman, Kreiman, Miller, & Poggio, 2008).

An interesting question that arises from our results is whether the online maintenance of temporal patterns is functionally relevant for the successful formation of memories. We could demonstrate subsequent memory effects for power decreases here, because a minimum of 15 trials per condition can yield stable power estimates (Hanslmayr, Spitzer, & Bäuml, 2009). We could further link power decreases to the presence of content specific temporal patterns; however because the trial count of forgotten associations for most of the subjects was too low for stable similarity estimates, it is not clear whether these patterns are functionally involved in memory formation. Specifically, the present study was designed to produce a sufficient number of remembered trials and we consequently could not contrast stimulus-specific temporal patterns between remembered and forgotten associations. Repeating this study in a longer and more adaptive design, could therefore allow for the contrast of pattern maintenance during successful and unsuccessful memory formation.

Additionally, future studies should address whether content-specific temporal patterns are causally involved in memory formation, either by disrupting content specific temporal patterns and therefore tampering with memory formation or even by artificially introducing spurious patterns to cause forged associations.

